# Modeling Global Genomic Instability in Chronic Myeloid Leukemia (CML) using patient-derived induced pluripotent stem cells (iPSC)

**DOI:** 10.1101/2022.09.09.505857

**Authors:** Gladys Telliam, Christophe Desterke, Jusuf Imeri, Radhia M’Kacher, Noufissa Oudrhiri, Estelle Balducci, Micheline Fontaine-Arnoux, Hervé Acloque, Annelise Bennaceur-Griscelli, Ali G. Turhan

## Abstract

Using induced Pluripotent Stem Cell (iPSC) technology, it is now possible to reprogram the primary leukemic cells to pluripotency and generate a major source of stem cells. To determine the feasibility of modeling global genomic instability characterizing chronic myeloid leukemia (CML) progression towards blast crisis (BC), we have used a patient-specific iPSC line that has been submitted to in vitro mutagenesis with the mutagenic agent N-ethyl-N-nitrosourea (ENU). Increased genomic instability was validated using g-H2AX and micronuclei assays. As compared to non-mutagenized cultures, these iPSC generated an increased numbers of progenitors (x 5-Fold) which proliferated in liquid cultures with a blast cell morphology. CGH array analyses were performed using ENU-treated iPSC-derived hematopoietic cells in two different time points as compared to leukemic iPSC cultured without ENU. Several cancer genes were found to be involved in the ENU-treated cultures, some known to be altered in leukemia (BLM, IKZF1, NCOA2, ALK, EP300, ERG, MKL1, PHF6 and TET1). Remarkably, transcriptome geodataset GSE4170 (Radich et al., 2006) allowed us to associate 125 of 249 of the aberrations that we detected in CML iPSC, with genes already described during progression of CML to BC (p-value =9.43exp32, after ANOVA with 1000 permutations). 38 most predictive aberrations allowed perfect reclassification of BC and chronic phase samples by unsupervised classification. Among these candidates, eleven of them have been described in CML physiopathology and connected to tyrosine kinase inhibitor (TKI) resistance and genomic instability. Overall, these results demonstrated that we have generated for the first time to our knowledge, an in vitro genetic instability model in CML, reproducing genomic events described in patients with BC. This tool using patient-derived iPSC could be of interest to study CML progression, to discover novel targets and eventually to test novel therapies.

## INTRODUCTION

Chronic myeloid leukemia (CML) is the prototype of a clonal malignancy of the hematopoietic stem cell (Sloma et al., 2010). The initiation of the disease by the appearance of Ph1 chromosome in a very primitive hematopoietic stem cell is followed invariably by the progression of the clone by acquisition of novel abnormalities leading to clonal progression, accelerated and blast phases of the disease (Kantarjian, 1987). This natural history has now been modified by the use of tyrosine kinase inhibitors (TKI) as first line therapies ensuring, in patients with deep molecular response, prolongation of their survival on therapy (O’Brien et al., 2003). However, resistances to several lines of TKI occur and accelerated and blast phases are observed potentially dues to clonal selection by TKI therapies (Shah et al., 2002; Soverini et al., 2006). Seminal and now classical work have established using serially collected patient samples, main cytogenetic and molecular events observed in patients during the progression of the disease towards accelerated and eventually blast phase. These include cytogenetic abnormalities identified as “major” and “minor” routes (Crisan et al., 2015; Mitelman, 1993). Several work performed using patients samples and models have also identified during the last two decades major molecular events involved in this progression such as, TP53 mutations (Perrotti et al., 2010). BCR-ABL expressing cells are also prone to develop a mutator phenotype (Canitrot et al., 1999) which could be due to the presence of an increase oxydative stress (Koptyra et al., 2008) which lead to clonal selection under the influence of TKI therapies (Liu and Wang, 2015). CML cells are finally doomed by the deficiency of several DNA repair mechanisms involving DNA-PKcs (Deutsch et al., 2001) BRCA1 (Deutsch et al., 2003) as well as abnormalities of specific DNA Repair genes such as Bloom (Slupianek et al., 2005) and other repair mechanisms such as NER (Canitrot, 2003) MMR (Stoklosa et al., 2008). Blast crisis is defined by a differentiation arrest decision of the leukemic progenitors and insensitivity to TKI. This alteration of the differentiation is due to this accumulation of genetic abnormalities.

iPSC technology offers the possibility of obtaining unlimited numbers of pluripotent cells which, despite difficulties of inducing definitive haematopoiesis (Lerou and Daley, 2005), can be used for modelling some aspect of hematopoiesis, as it is possible to generate terminally differentiated progenitors and differentiated cells. Blast crisis is characterized by the progressive occurrence of genomic abnormalities dues to BCR-ABL-induced genetic instability and (Chereda and Melo, 2015). CML patient-derived iPSC express BCR-ABL (Telliam et al 2016, Telliam et al 2022) and can allow the possibility of inducing global abnormalities by in vitro mutagenesis. In order to explore the possibility of generating such model from an iPSC derived from a CP-CML patient, we have chosen an in vitro mutagenesis approach using N-ethyl-N-nitrosourea (ENU). Here we show that CML patient derived IPSC can be used to generate an experimental model mimicking cytological and genomic instability patterns observed in primary blast crisis cells. Moreover, we show these abnormalities induced by this technology are representative of those described in large database of genomic events described in primary CML patients in blast crisis.

## MATERIALS AND METHODS

### Patient-derived iPSC and characterization

Generation of CML iPSC PB32 has been previously described (Telliam et al., 2016). Briefly, peripheral blood cells were obtained at diagnosis from a 14-year old CML patient with informed consents according to the declaration of Helsinki. This patient was treated with Imatinib mesylate as a first line therapy. At 12 months post-therapy there was no major molecular response and an allogenic bone marrow transplant was performed. The patient remains in complete remission since the transplant performed in 2004. To generate iPSC, cryopreserved CD34+ cells were used as previously described (Telliam et al., 2016). All analyses were performed using a polyclonal stock of iPSC. Control cells used for these experiments included PB33, an iPSC cell line that was obtained from bone marrow CD34+ cells using Sendai virus. CML and control iPSC have been characterized for their pluripotency using cell surface markers as well as by generation of in vivo teratoma assays in immunodeficient mice (Telliam et al., 2016).

### iPSC and ESC cultures

hIPS cell lines were maintained on the mitomycin-C-inactivated mouse embryonic fibroblast feeder cells with DMEMF12 supplemented with 0.1mg/ml bFGF, with or without N-ethyl-N-nitrourea 10μg/ml or 50μg/ml and treated 2h with 1mg/ml Collagenase type IV (Gibco by life technologies ref 17104-019).

### Embryoid body assays

To induce embryoid body (EB) formation, ES and iPS cells at day 6–7 after cell passage, were treated with collagenase IV. Clumps were cultured in Iscove’s modified Dulbecco’s medium (IMDM, Invitrogen) supplemented with 1% penicillin/streptomycin, 1 mM L-glutamine, 15% fetal calf serum (FCS, Invitrogen), 450 μM monothioglycerol, 50 μg/mL ascorbic acid (Sigma Aldrich) and 200 μg/L transferrin (Sigma Aldrich) and supplemented with hematopoietic cytokines: 100 ng/mL stem cell factor (SCF), 100 ng/mL fms-like tyrosine kinase 3 ligand (Flt-3L) and 50 ng/mL thrombopoietin (TPO) (all from Peprotech). ES and IPS derived EB were cultured in ultra-low attachment 6-well plates (Costar) for 16 days. Media was changed two or three times depending on EB proliferation. All cultures were incubated at 37°C in 5% CO2

### Blast-colony forming assays

Blast colony forming cell (Bl-CFC) assays were performed using iPSC cell pellets resuspended in Stemline II Hematopoietic Stem Cell Expansion medium (Sigma-Aldrich S0192) supplemented with 1% penistreptomycin and L-glutamine, 50ng/ml rhVEGF (Peprotech) and 50ng/ml rhBMP4 (Peprotech). After 48h we added 20ng/ml of rhSCF (Peprotech), rhTPO (Miltenyi Biotec) and rhFLT3L (Peprotech). The embryoid bodies at day 3.5 were collected, dissociated with pre-heated stable trypsin replacement enzyme TrypLE Express (gibco by life technologies ref 12605-10) and filtered with 40μm Nylon Mesh sterile cell strainer (Fisher Scientific). Cells were resuspended in Stemline II Hematopoietic Stem Cell Expansion medium (Sigma-Aldrich S0192) supplemented with 1% peni-streptomycin and L-glutamine, 50ng/ml rhVEGF, rhBMP4, rhFLT3L and rhTPO, 20ng/ml FGFb (Peprotech) and 5Units/ml EPO (Peprotech) and transferred in Methocult SF H4436 (Stem cell technologies) for 5 to 7 days.

### Hematopoietic differentiation

CFC assays from ES and iPS cells from EB or BL-CFC were plated at 10×10^3 cells/mL into MethoCult GF (H4435 StemCell Technologies). Hematopoietic CFC was counted at day 19.

### Western Blots

Cells were lysed in ice with RIPA buffer containing (NaCl [200mM], Tris [pH 8; 50mM], Nonidet P40 [1%], acide deoxycholate [0.5%], SDS [0.05%], EDTA [2mM]) supplemented with 100μM phenylmethylsulfonyl fluoride (PMSF), 1mM sodium fluoride (NaF), 1mM orthovanadate (Na3VO4). Separation of proteins was done by electrophoretic migration on a 3-8% polyacrylamide gel under denaturing conditions. The proteins were then transferred in a semi-liquid condition on PVDF membrane pre-activated in methanol. After saturation with TBS Tween 5% BSA for 1h and hybridization of the membranes with primary (Anti-phosphoHistoneH2A.X (Ser139) clone JBW Millipore) and secondary antibodies coupled to HRP. Membranes were revealed by chemiluminescence with SuperSignal West Dura or Femto reagents and data acquired using G:BOX iChemi Chemiluminescence Image Capture system.

### Evaluation of hematopoietic cells phenotypes

At day + 19 of cultures, the methylcellulose was washed with PBS. Living cells were counted in trypan blue and stained with the following antibodies in PBS 4%BSA at 4°C 45 min: CD45Pe-Vio770 (Miltenyi), CD34Vioblue (Miltenyi), CD41aPE (BD), CD43FITC (Miltenyi), CD31FITC (Beckman Coulter), CD235aAPC (BD), IL1-RAP APC (R&D), CD38PE (Miltenyi), CD71PE (BD), CD14FITC (Beckman Coulter), CD33APC (BD), CD133APC (BD), BB9PE (BD), SSEA1PE (BD). After 1h, cells were washed and resuspended in PBS 4%BSA with viability staining reagent 1μg/ml (7-AAD) 7-aminoactinomycin D (Sigma Aldrich). Stained cells were analyzed with a MACSQuant 10 (Miltenyi Biotec) flow cytometer and Flowjo analysis software.

### ENU experiments

CML iPSC and control iPSC cells were cultured in the presence or in the absence of ENU at the concentration of 10 μg/ml with daily addition of ENU as described above. Experiments performed used iPSC cultured in ENU for either 40 days or 61 days. CGH array experiments were performed using EB’s derived hematopoietic cells and Bl-CFC generated from ENU-treated cultures as compared to cultures not treated with ENU.

For kinetic analysis, we recovered IPSC colonies on mouse embryonic fibroblasts (MEF). The “clumps” of colonies have been incubated in DMEMF12 medium supplemented with 50μg/ml ENU at 37 ° for 2, 10 and 30 min. After treatment, the protein pellet was extracted from each condition in order to perform a kinetic analysis of the level of γH2AX by Western blot.

### Cytogenetic analyses

Caryotype analyses were performed using cell pellets collected at different time points using standard methods as previously described (Telliam et al., 2016).

### Micronuclei analyses

The presence of micronuclei and anaphase bridges before and after exposure to ENU were analyzed as described previously [Zaguia et al 2020]. Briefly, cells were incubated in a humidified atmosphere of 5% (v/v) CO2 in air at 37 °C until arrest and cell spread. The culture medium was discarded and hypotonic shock was induced by incubating the cells with 18mM KCl at room temperature. The cells were then fixed with acetic acid/ethanol (1:3, v/v), and the cell suspension was dropped onto the slides. The slides were stored at −20 °C until further use. Telomere and centromere staining was performed in order to detect the nature of micronuclei: with only telomere staining resulting mainly from acentric chromsomes or with telomere and centromere staining which predominatly contained lagging chromosomes.

Automatic scoring of MN was performed using MNScore software (version 3.8.101 MetaSystems, Althaussen, Germany) with a Metafer 4 image analyser (MetaSystems, Althaussen, Germany) comprised of a Zeiss Axioplan 2 imager to detect MN. An operator validated and excluded the false MN in cells. For each sample, 1000 cells were scored.

### DNA Extraction

iPSC, EB’s and Bl-CFC obtained from either ENU-treated or not treated CML iPSC as well as from control iPSC were used for DNA extraction using standard methods with DNeasy Kit (Qiagen).

### Array Comparative Genomic Hybridization

To perform CGH arrays, we have used genomic DNA from ENU-treated iPSC derived EB’s and Bl-CFC’s compared to CML iPSC without ENU treatment. We followed Agilent Oligonucleotide Array-Based CGH for Genomic DNA Analysis protocol. Briefly, we performed digestion from 500ng of DNA with SureTag DNA labeling kit (4-packs). After gDNA amplification CML iPSC was labeled with Cyanine 3 and the tested samples with Cyanine 5. Assembled chambers were loaded into the oven rotator rack, hybridization were performed at 67°C for 24 hours at 20 rpm.

### Transcriptome dataset

Rosetta / Merck Human 25k v2.2.1 microarray Array data matrix of normalized log ratios from GSE4170 was downloaded on Gene Expression Omnibus website: (http://www.ncbi.nlm.nih.gov/geo/query/acc.cgi?acc=GSE4170), and annotated with annotation plateform GPL2029 (http://www.ncbi.nlm.nih.gov/geo/query/acc.cgi?acc=GPL2029). In this dataset CD34+ cell samples belonging to the 3 experimental groups corresponding to the different evolution phases of chronic myeloid leukemia were analyzed: chronic phase, accelerate phase and blastic crisis were analyzed (Radich et al., 2006)

### Bioinformatics

Boxplot, two-sided Student t.test with Welch correction were performed in R Software version 3.2.3. Heatplot was performed with made4 package (Culhane et al., 2005). Functional enrichment was performed with Go-Elite Standalone software version 1.2 on the Gene Ontology Biological Process, KEGG and CommonsPathway databases included in Homo sapiens EnsMart77Plus (Ensembl – Biomart) update (Zambon et al., 2012). Functional interaction networks were performed with Cytoscape software version 3.2.1 (Cline et al., 2007). Unsupervised principal component analysis was performed on gene expression profile with R package FactoMiner; p-value was calculated by group discrimination on first principal component axis. Pubmed gene priorization was realized with gene valorization web application (Brancotte et al., 2011). Circosplot on contingency table of Pubmed gene priorization was realized with Circlize R package (Gu et al., 2014). Circosplot of genomic aberrations with HG19 coordinates were performed with OmicCircos R package (Hu et al., 2014). ROC curves were performed with ROCR R package. Genomic aberration gene candidates were also matched with important databases such as: database PLURINET for genes implicated in pluripotency (Müller et al., 2008), cancer genes COSMIC database: Catalogue of somatic mutations in Cancer (http://cancer.sanger.ac.uk/census) (Futreal et al., 2004) and transcription factors (Vaquerizas et al., 2009)

## RESULTS

### Long term ENU exposure induces an enhancement of CML-IPSC derived hematopoiesis

To this purpose we compared EB or Bl-CFC and hematopoietic cells from CFC assay derived from IPSC cultured under normal culture conditions or after ENU exposure (**figure 1B**). As we can see figure 1B, after 2 months of ENU exposure CML-IPSC showed a significant increase of hematopoietic progenitor potential (Mann-Whitney p= 0.0006) (**figure 1B**). Interestingly, in CFC assays generated from PB32-ENU cells we have observed larger colonies which proliferated in methycellulose cultures (**Figure 1B**) as well as in liquid cultures in the presence of hematopoietic growth factors (SCF, TPO, Flt3L). Cytological features of these cells using May-Grunwald stain revealed the presence of myeloid cells with precursor and mature (myelocytes, metamyelocytes, neutrophils, monocytes/macrophages) cells as well as cells with undifferentiated blast cell morphology reminiscent of blast cells (**Figure 2B**). FACS analysis of these cells showed that the majority of cells expressing CD45, with positivity of CD31, CD38, CD43 markers (**Figure 3**). Hemangioblast marker BB9, as well as CD133, CD26, SSEA1, CD13, CD33 and CD71-expressing cells were also detected (**Figure 3**).

**Figure 1:**
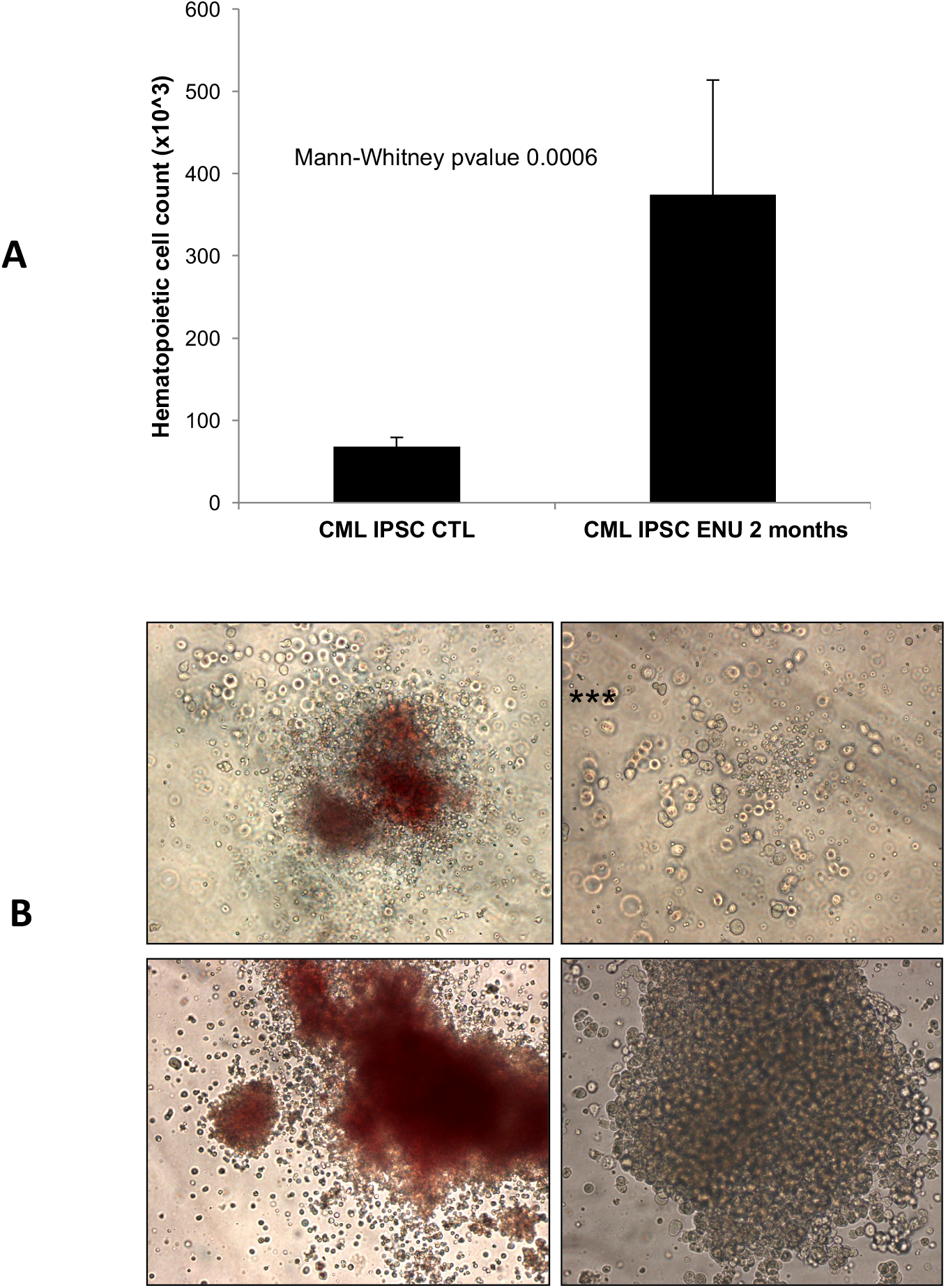
Increased hematopoietic potential in CML-iPSC under ENU exposure. **(A)** Hematopoietic progenitor count from CML IPSC before or after exposure to 10μg/ml ENU during 2 months. Statistical Mann-Withney t.test was performed with PRISM software. **(B)** CFC generated in methycellulose cultures revealing large colonies with myeloid differentiation features associated with blast colonies.

**Figure 2:**
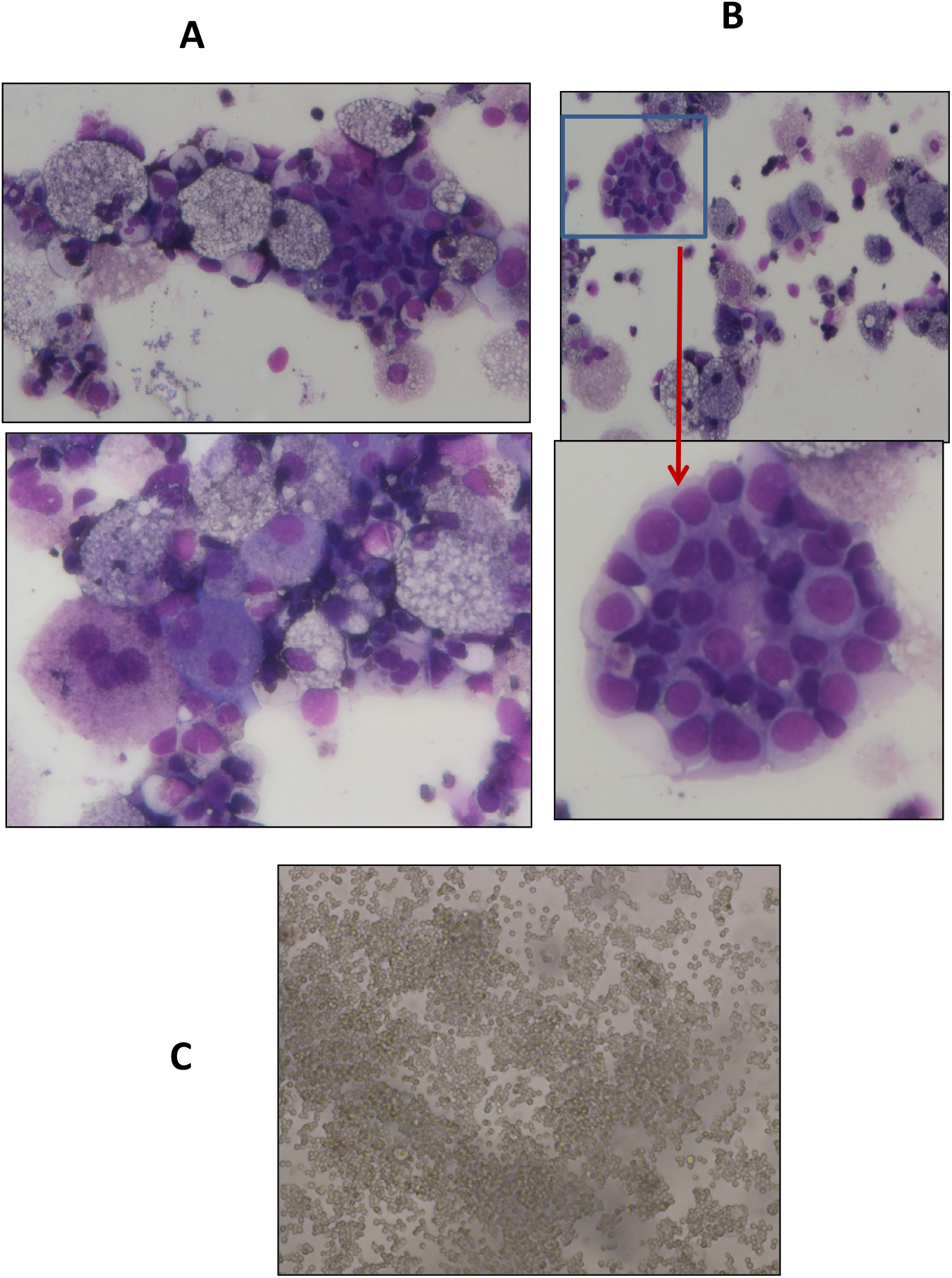
May-Grunewald staining of hematopoietic colonies. **(A)** Evidence of myeloid differentiation along with **(B)** blast colonies. **(C)** Proliferation of blast cells in liquid cultures in the presence of SCF, TPO and Flt3-ligand for > 2 weeks.

**Figure 3:**
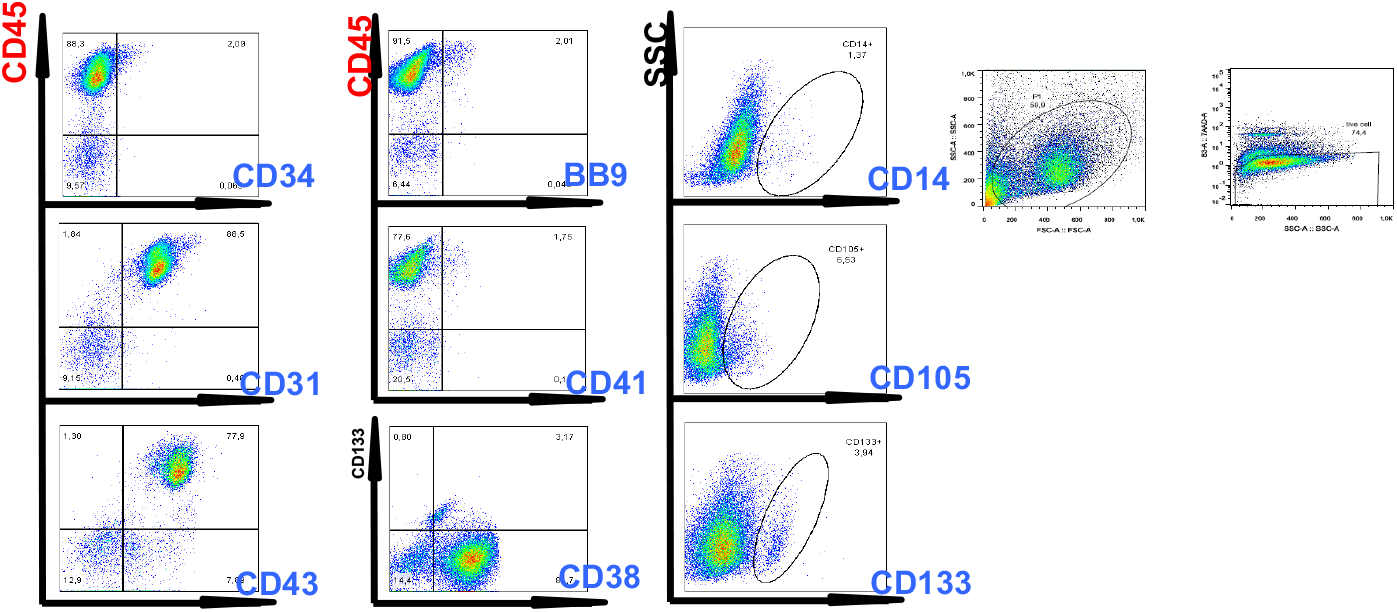
Flow cytometry evaluation of hematopoietic cells generated after **from ENU-treated** CML iPSC. As can be seen in this Figure, double staining experiments showed the presence of CD45 + / CD43+ and CD45+/CD38+ cells demonstrating generation of hematopoietic cells with primitive characteristics.

### Genomic instability of CML-IPSC induced by BCR-ABL under ENU exposure

Differences observed in the behavior of PB32 and PB32-ENU cells suggested that ENU could enhance the DNA repair abnormalities known to be inherent to CML cells. We have performed for this purpose a short-term exposure to ENU of our CML iPSC and analyzed the kinetics of accumulation of phospho-γH2AX in these cells.

We cultured IPSC colonies on mouse embryonic fibroblasts (MEF). The “clumps” of colonies have been incubated with ENU at 37 ° for 2, 10 and 30 min. After treatment, the protein pellet was extracted from each condition in order to perform a kinetic analysis of the level of γH2AX (**Figure 4A**). Kinetic analysis of an IPSC control cell line revealed an increase of γH2AX expression level at 10 minutes of treatment with ENU with a progressive decrease over time. These observations suggest that ENU induces time dependent DNA breaks and the DNA repair system is set up around 10 minutes after detection of γH2AX signal. However, CML-IPSC cell line showed a gradual accumulation of γH2AX (**figure 4B**) with a significant increase after 10 min and persistence of the phosphorylation after 30 min of ENU treatment (n=3, Two way ANOVA with Bonferroni’s correction) (**figure 4C**). These results suggested the presence of a baseline increase of double strand breaks in CML iPSC which persisted after ENU exposure due potentially to a deficient DNA repair system.

**Figure 4:**
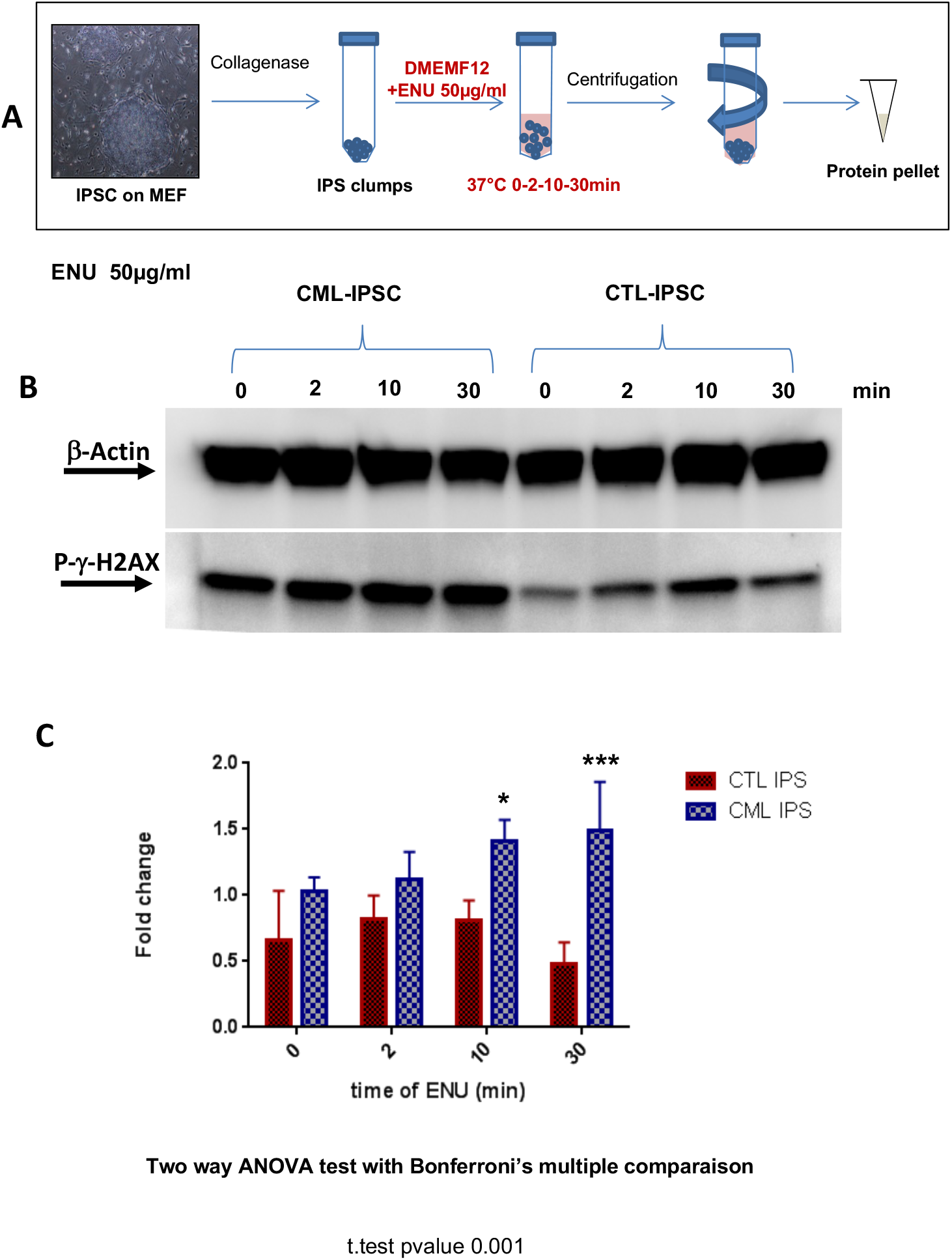
Evaluation of the phosphorylation of γ-H2AX in the CML iPSC after short-time ENU exposure. **A-** CML iPSC cultured on murine embryonic fibroblasts were collected after collagenase treatment and the pellets subjected to a short time exposure to ENU followed by protein extractions. **(B)** Western blot analysis of the phosphorylation level of the histone variant gamma-H2AX (γH2AX) on Ser-139 after exposure of the CML-IPS cell line and control iPSC to ENU at 50μg/ml for 0, 2, 10 and 30 minutes (30μg of protein per conditions). **(C)** Quantification of the γH2AX as compared to ß-Actin with ImageJ software, histogrammes represent means of 3 independent experiments. A Two-way ANOVA test with multiple comparison and Bonferroni’s correction was made with PRISM software.

The potential genomic instability of PB32-ENU cells was also evaluated by the micronucleus assays (Zaguia et al 2020) which is a recognized marker of the chromosomal instability of the eukaryotic cells. As can be seen in the Supplementary Figure 3, the presence of spontaneous micronuclei was significantly higher in the mutagenized CML iPSC lines (PB32-ENU) as compared to the non-mutagenized counterpart and to the control cell line (PB33-passage 53), validating the genomic instability of the mutagenized CML-iPSC (**Supplementary Fig 3**).

We then wished to evaluate the presence of other features of cytogenetic and genomic abnormalities in PB32-ENU cells as compared to PB32 not treated with ENU. To this purpose, we have performed cytogenetic analyses on CML iPSC after long-term exposure to ENU (PB32-ENU) as compared to PB32 cultured without ENU. As can be seen in Table 1, several cytogenetic abnormalities were identified in addition to t(9;22) (**Table 1**). Amongst these abnormalities (64 mitosis analyzed), we have noted the deletion of der 9q (3 mitosis / 64) (5%) deletion of Chromosome 21 (5 mitosis / 64) (8%) (**Table 1 and Supplementary Figure 1**). Despite several passages, control iPSC cells exposed to ENU (PB33) did not develop any detectable cytogenetic changes (Suppl Figure 2). To determine the genomic events at the molecular level, we have performed CGH array experiments by comparing EB derived CFC and Bl-CFC DNA generated from PB32-ENU as compared to PB32 cultured without ENU.

**Table 1.** Evaluation of Additional Chromosomal Abnormalities in iPSCs before and After ENU-induced mutagenesis

| **IPSC Line** | **# of Chromosome Involved** | **Mitosis Ratio** | **%** |
| --- | --- | --- | --- |
| **PB32 + ENU (Ph1+)** | 21 | 5 loss / 64 | 8 |
|  | 9 | 3 loss / 64 | 5 |
|  | 17 | 1 loss / 64 | >2 |
|  | 19 | 1 loss / 64 | >2 |
|  | 20 | 1 loss / 64 | >2 |
|  | 13 | 1 loss / 64 | >2 |
|  | 1 | 1 loss / 64 | >2 |
|  | 2 | 1 loss / 64 | >2 |
| **PB32 without ENU (Ph1+)** | None | / | / |
| **PB33 + ENU (Ph1-neg)** | None | / | / |
| **PB33 without ENU (Ph1-neg)** | None | / | / |

PB32+ ENU : Ph1+ CML-IPSC line cultured with ENU
PB32 without ENU : Ph1+ CML-iPSC without ENU-mutagenesis
PB33+ENU: Control Ph1-negative cell cultured with ENU
PB33-Ph1- : Control Ph1-negative cell line

### CGH array analysis of PB32-ENU-derived hematopoietic cells

Figure 5 shows the experimental strategy to perform this analysis. As can be seen in this Figure 5, we have extracted DNA from either PB32-ENU EB-derived CFC or Bl-CFC and compared the genomic abnormalities to that of PB32 cultured all along without ENU (**figure 5**). Cytogenomics software from Agilent technologies with mosaic workflow allowed to detect 20 quantitative genomics aberrations: copy number variations CNVs which comprised 313 gene loci. After filtration on European Caucasian genomic polymorphism database still 295 gene loci are comprised in these genomic aberrations (**figure 6A and supplementary table 1**). Majority of the gene loci events are affected by gain of genomic DNA (69%) versus loss (31%) (**Figure 6B and supplementary table 1**). Matching these genomic aberrations with transcription factor database, cancer gene database and pluripotency gene database allowed to observed that these important deregulated actors are principally affected on chromosomes 5, 7, 21, 22, and X (**figure 6B**): majority of these genomic events implicated are transcription factors such as: zinc finger ZNF and IRX (**figure 6B**). Some pluripotency genes were also identified as well as cancer genes such as TET1, ALK, EP300, ERG, MKL1, PHF6 already described as been involved in leukemia. Functional enrichment of genomic alterations of EB-derived CFC allowed to highlight perturbations in megakaryocytic development with HBD, HBE1, HBG1 and HBB molecules and HDAC enzymes including EP300 and SIRT1 (**figure 6C and D**). Functional enrichment on Gene Ontology database allowed to identified SIRT1, EP300 and CDH13 as central molecules especially SIRT1 which is implicated in p53 dependent-DNA damage in response to hydrogen peroxide and cellular response to hypoxia. SIRT1 is also involved in negative regulation of NK-kappaB (**figure 6E and 6F**).

**Figure 5:**
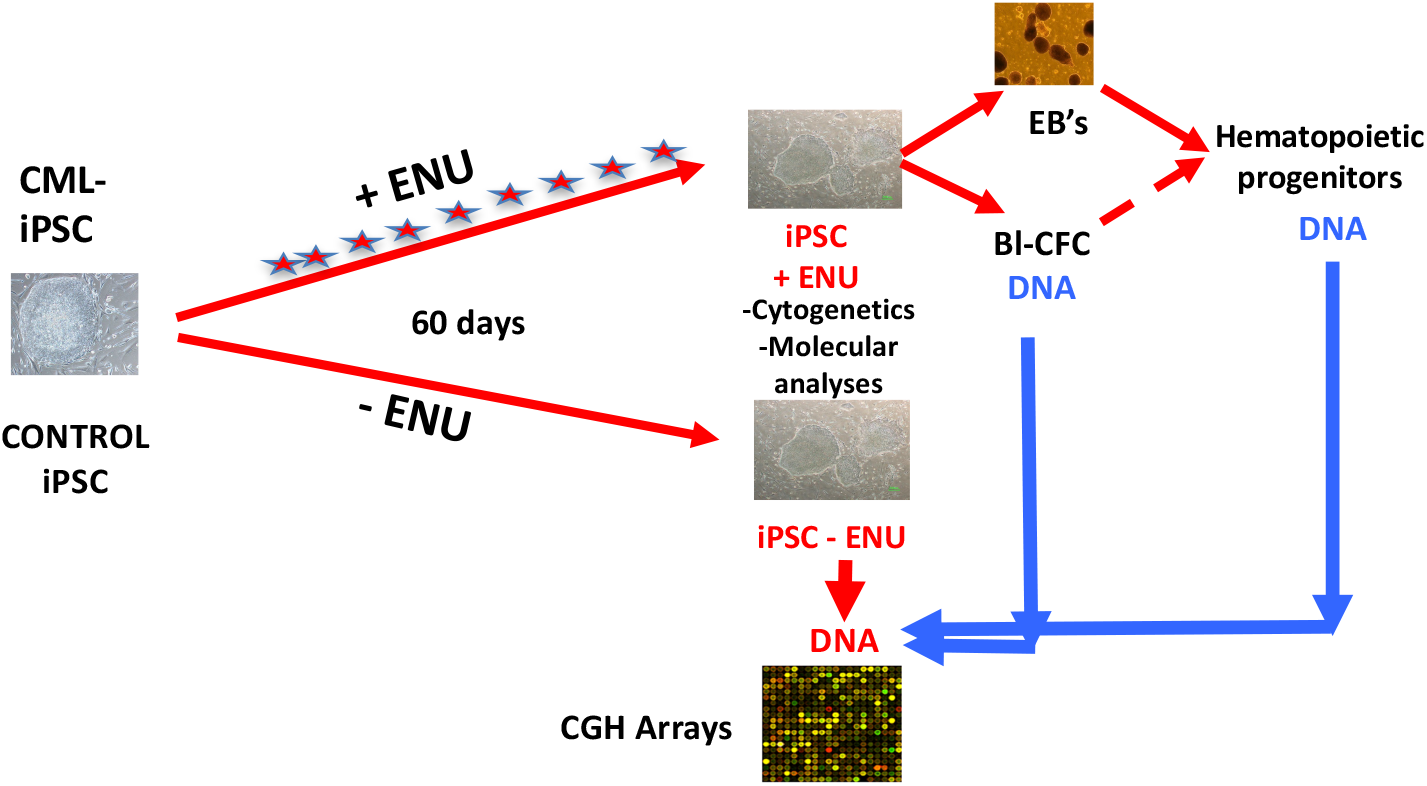
Experimental protocol used for aCGH experimental procedure. CML and control iPSC were cultured in murine embryonic fibroblast layers in the presence or in the absence of ENU for 60 days, with daily medium changes containing or not ENU. At day +60, DNA extracted from hematopoietic progenitors generated via EB’s or Blast-CFC’s were used in CHG arrays experiments using DNA extracted from iPSC cells without ENU exposure.

**Figure 6:**
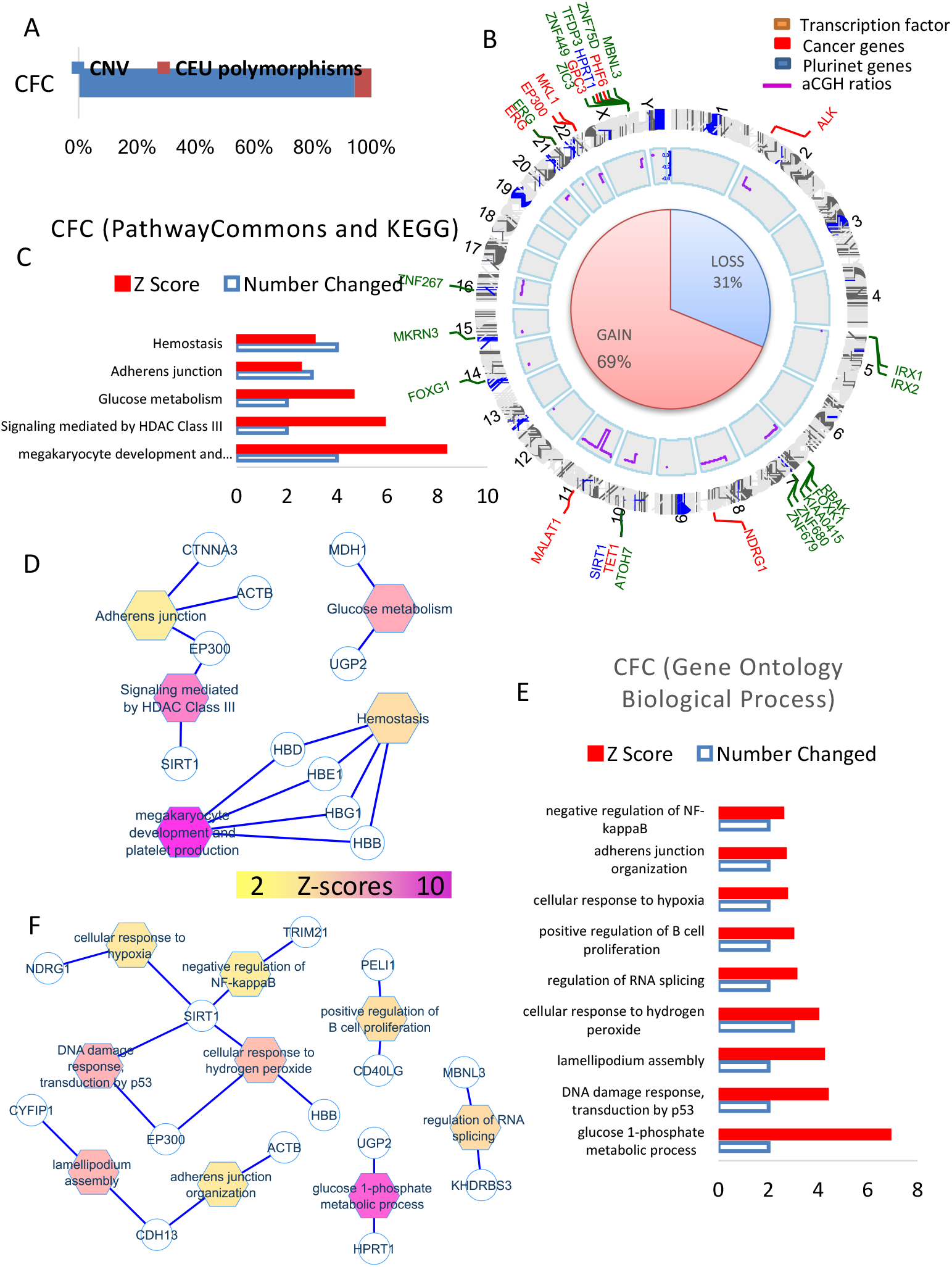
aCGH analysis of EB-derived CFC cells. **(A)** Quantitative genomics aberrations: copy number variations CNVs, **(B)** Matching of genomic aberrations with transcription factor database, cancer gene database and pluripotency gene database, circosplot show in green: 5 zinc finger ZNF genes and 2 IRX, in blue: pluripotency genes, in red: cancer genes. **(C-D)** Functional enrichment of genomic alterations of CFC on KEGG database. **(E-F)** Functional enrichment on Gene Ontology database.

When the same analysis was performed using DNA of Bl-CFC, 22 quantitative genomics aberrations were detected. These included copy number variations (CNVs) which comprised 332 gene loci. After filtration on European Caucasian genomic polymorphism database still 255 gene loci were found to be comprised in these genomic aberrations (**figure 7A and supplementary table 2**). Majority of the gene loci events represented loss of genomic DNA (71%) with loss of heterozygosity (23%) and 6% gains of DNA (**Figure 7B and supplementary table 2**). Matching these genomic aberrations with transcription factor database, cancer gene database and pluripotency gene database allowed to demonstrate that these important deregulated actors were principally involved on chromosomes 7, 8, 15, Y, and X (**figure 7B**): circosplot analysis also allowed to show that majority of these events involved comprised transcription factors such as MESP implicated in mesodermal cell migration and IKZF1 (**figure 7B and 7F**). Amongst the, few pluripotency and cancer genes we identified IDH2, NCOA2, IKZF1, BLM described as being involved in leukemia (**Figure 7B, 7F**). Functional enrichment of genomic alterations of CB32 IPSC on KEGG database allowed to highlight perturbations in hematopoietic lineage and cytokine-receptor interaction, affecting: TPO, CSF2RA, ILRA, PIK3RA, CRFL2 so cytokines and receptors allowing activation JAK-STAT pathway (**figure 7C and 7D**). Functional enrichment on Gene Ontology database allowed important implication of developmental molecules and especially sharing functionalities in mesodermal tissue (**figure 7E and 7F**) such as IKZF1 a transcription factor of the mesodermal development and MESP1 et 2 two transcription factors implicated in migration of mesodermal cells; also important implication of molecules in relation with the extracellular matrix, cell fate specification and DNA repair.

**Figure 7:**
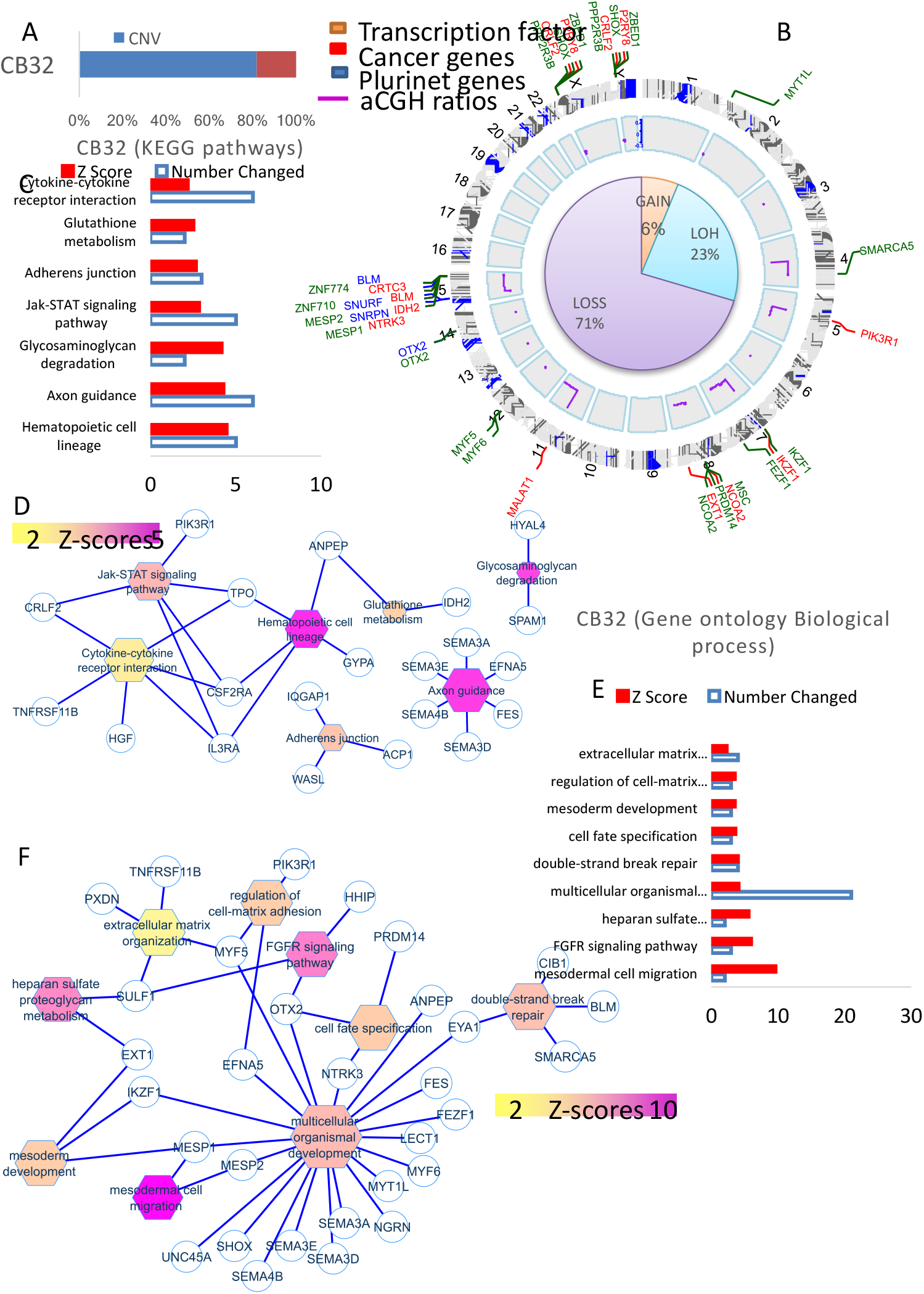
aCGH analysis of Bl-CFC progenitors cells. **(A)** Cytogenomics software from Agilent technologies with mosaic workflow: quantitative genomics aberrations (copy number variations CNVs which comprised 332 gene locus). After filtration on European Caucasian genomic polymorphism database 255 gene locus are comprised in these genomic aberrations. **(B)** Pourcentage of genomic loss, gains and LOH (Loss of heterozygosity), match of genomic aberrations with transcription factor database, cancer gene database and pluripotency gene database. **(C-D)** Functional enrichment of genomic alterations of Bl-CFC on KEGG database. **(E-F)** Functional enrichment on Gene Ontology database.

### Comparison of genomic aberrations identified by aCGH in CML iPSC as compared to primary leukemic blast crisis gene profiling

Transcriptome of CML CD34+ cells in the 3 different phases of the disease: chronic phase (CP), accelerated phase (AP) and blast crisis (BC) were analyzed to extract information of genomic aberrations that we observed in CML PB32-ENU_derived hematopoietic progenitors. On 249 candidate genes, a supervised one-way ANOVA was performed between the 3 phases of the disease (with 1000 permutations and p-value threshold less 1E-3): 125 genes were found significant and unsupervised principal component analyses performed with them allowed to have a very significant group discrimination (p-value of discrimination 9.43E-32, **figure 8A and supplementary table 3**). Interestingly, among these genes some were identified EB-derived CFC’s and some in Bl-CFC-derived cells (**Figure 8B**). Unsupervised classification performed with the gene candidates mostly correlated to the progression of the disease, allowed to obtain a perfect reclassification of the chronic phase samples as compared to the blast crisis samples (**figure 8C**). NCBI gene valorization performed one these best candidates correlated to the progression disease allowed to identified eleven genes (**figure 8D**) which are well known in CML pathology, genomic instability and tyrosine kinase inhibitors related keywords in published CML literature (**Figure 8D**). Majority of these candidates including a group of 7 genes (7 of 11) are predictive for the chronic phase (**figure 8D**), whereas the other group are predictive for blast crisis (4 of 11) (**figure 8D**). Among genes candidates not reported in CML pathophysiology, were found to be predictive of the blast crisis, such as DDX family such as DDX50 and DDX21, very well known in DNA repair and the implication of which is unknown in CML.

**Figure 8:**
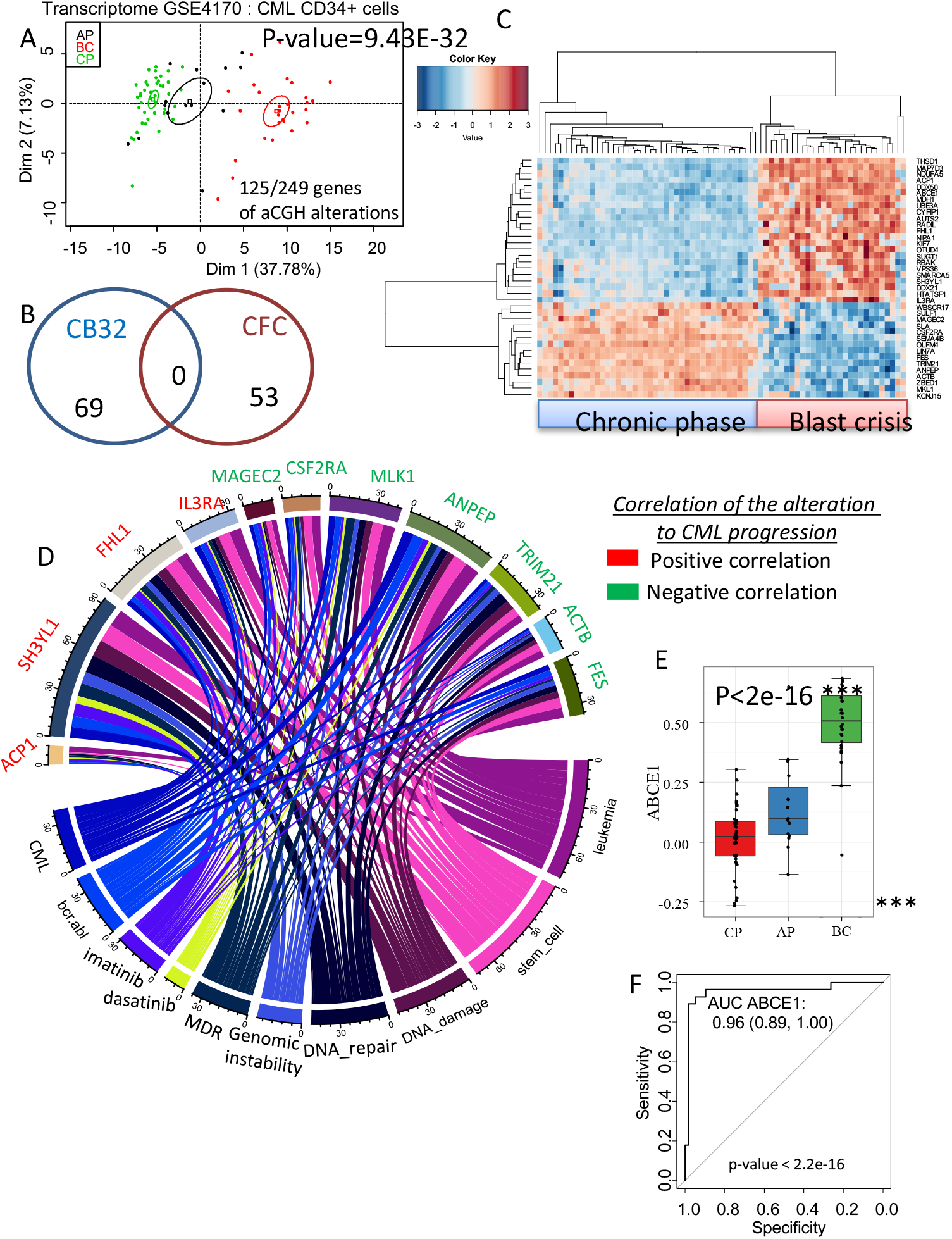
Comparative analysis of genomic aberrations discovered in CML-iPSc aCGH with the gene profiling described in primary CML CD34+ cells. **(A) In silico analysis of geniomic aberrations found in our study as compared to** gene profiling of CML CD34+ cells described in Radish ( ) in the 3 different phases of the disease: chronic phase (CP), accelerated phase (AP) and blast crisis (BC) were analyzed to extract information of genomic aberrations that we observed under ENU exposure, supervised with one-way ANOVA. **(B)** Genes related to CFC or Bl-CFC aCGH analysis. **(C)** Classification of the chronic phase samples as compared to the blast crisis samples. **(D)** NCBI gene valorization performed one the best candidates correlated to the progression disease and well known in CML pathology, genomic instability and tyrosine kinase inhibitors related keywords; in green: predictive candidates to chronic phase, in red: predictive candidates to blast crisis.

## DISCUSSION

In the current era of TKI therapies, the progression towards BC became rare but still occurs even after several years and correlates with resistance to several lines of therapies. Several CML cell lines have been generated from patients in established blats crisis, the first being K562 (Lozzio and Lozzio, 1975). Mechanisms underlying the progression of CML towards have been extensively studied during the last four decades (Perrotti et al., 2010) using serially collected patient samples or in vitro and in vivo models, essentially by analyzing blast crisis cell lines such as K562 or by overexpressing along with BCR-ABL, genes involved in blast crisis such as NUP98 (Sloma et al). Major cytogenetic events (Crisan et al., 2015) as well as molecular events contributing to the differentiation arrest have been identified (Melo and Barnes, 2007) (Perrotti et al, 2010). More recently, global genomic analyses performed in primary BC samples showed the enrichment of CML genome by the occurrence of mutations, essentially of the polycomb repressive complex (PRC) (Ko TK et al, Blood 2020). However, these tools do not allow to study progressive changes occurring during the clonal evolution of the disease, which require serial samplings. Similarly, blast crisis in CML can be modeled using murine CML models by using complementation assays using BCR-ABL and an oncogenic fusion protein such as NUP 98/ HoxA9 (Dash et al., 2002; Neering et al., 2007). Although these studies can be a major interest to study the in vivo behavior of cells, they are not appropriate to study human leukemia in the context of a given genomic background of a patient.

In this work we wished to evaluate the feasibility of generating a genetic instability model of CML using patient-specific induced pluripotent stem cells. An iPSC cell line has been established from a patient with TKI-resistant CML and previously has been characterized (Telliam et al., 2016). These cells exhibited all features of pluripotency including cell surface and in vivo teratoma generating abilities (Telliam et al., 2016) and have been cultured in a pluripotent state for several passages. We have shown that these cells express BCR-ABL and BCR-ABL-signaling which appears to be preserved during differentiation towards hematopoiesis generated via embryoid body or blast-cell colony assays (Telliam et al, Stem Journal, 2022). Using clonogenic cell assays, we have shown that this IPSC cell line PB32 exhibits an increased hematopoietic potential as compared to standard ES cells such as H1 or control iPSC without BCR-ABL expression (Telliam et al, 2016).

To mimic in vitro genetic instability, we have used ENU, a well-established mutagen and alkylating agent (Acevedo-Arozena et al., 2008). We have previously shown that ENU can be used in CML cell lines to generate ABL-kinase mutations that can be selected in vitro by TKI and allows in vitro production of clinically relevant mutations such as T315I substitution (Aggoune et al., 2014). After testing the appropriate dosing experiments, we have used ENU in vitro during each daily medium changes. PB32-ENU cells have thus been cultured for > 60 days with PB32 counterparts without ENU. This treatment did not appear to induce any toxicity and no morphological changes in iPSC (data not shown). However, in hematopoiesis induction experiments, we have observed a major increase of hematopoietic potential using either EB or Bl-CFC-derived hematopoietic cells (**Figure 1**). Hematopoietic cells generated in these cultures could be expanded in short term cultures, revealing the presence of myeloid precursors and blast-like cell reminiscent of an accelerated phase of CML (**Figure 2**) whereas such pattern was not observed in iPSC cultured without ENU. Interestingly, cytogenetic analyses performed in PB32-ENU cells revealed the presence of several chromosomal abnormalities in addition to Ph1 chromosome (Figure and Table 1). These included the presence, in several mitoses (3 / 64 analyzed) deletion of the chromosome derq9, which has been shown to occur at diagnosis (Kreil et al., 2007) but also during progression towards blast crisis (Huntly et al., 2002; Reid et al., 2002). Another significant abnormality observed was the deletion of chromosome 21 in several mitoses in PB32-ENU cells (**Table 1**). None of these abnormalities are representative of a major or minor route but they have been described in CML patients during blast crises (Storlazzi et al., 2002). The analyses of PB32 ENU cells as compared to PB32 revealed also the presence of extensive micronuclei formation, suggesting the induction of significant genomic instabilities in these cells (**Supplementary Figure 3**). Micronuclei are characteristics of cells undergoing DNA damage especially in malignant cells (Terradas et al., 2010). They are formed during mitosis and can be due to the presence of double strand DNA breaks (Terradas et al., 2010). Increased micronucleus frequency has been shown to be a feature of myelodysplastic syndromes and has been shown to be higher in patients with refractory anemia with excess of blasts (Kuramoto et al., 2002). To analyze the chromosomal abnormalities at the genomic level, we performed CGH array analyses to compare PB32-ENU-derived hematopoietic cells or Bl-CFC’s as compared to PB32 without ENU treatment (**Figure 5**). This analysis revealed several major genomic aberrations (**Figure 6 and 7**). As can be seen in figure 6, in hematopoietic cells we have observed several losses (31%) or gains (69%) interesting all chromosomes and some of major genes involved in oncogenesis such as TET1, ALK, ERG and MALAT1 (**Figure 6**). Several transcription factor genes were also involved such as ZNF or ERG family of transcription factors (**Figure 6**). Functional enrichment on Gene Ontology database showed the implication of SIRT1, EP300 and CDH13, SIRT1 being involved in TP53-dependent DNA damage.

The analysis performed using comparative hybridization of Bl-CFC DNA, yielded additional information with losses (71%) and gains (6%), and matching these genomic aberrations with transcription factor database, cancer gene database and pluripotency gene database allowed to confirm that these affecting essentially chromosomes 7, 8, 15, Y, and X (**figure 7B**). Circosplot analysis showed that the majority of these abnormalities implicate transcription factors such as MESP (implicated in mesodermal cell migration) and IKZF1 (**7B and 7F**). Amongst pluripotency genes, we have identified cancer genes (IDH2, NCOA2, IKZF1, BLM) which are already described as been involved in leukemia. Functional enrichment of genomic alterations of CB32 IPSC on KEGG database allowed to highlight perturbations in hematopoietic lineage and cytokine-receptor interaction, affecting: TPO, CSF2RA, ILRA, PIK3RA, CRFL2, cytokines and receptors allowing activation of JAK-STAT pathway (**figure 7C and 7D**).

We then asked whether the abnormalities identified using CGH arrays could be matched with genomic aberrations described in primary leukemic cells from blast crisis patients. Amongst 249 genes identified by aCGH in CML iPSC, 125 have been found to be already described in CML progression (**Figure 8 and Supplementary Table 3**). These positively correlating genes include IL3RA, ACP1, FHL1, SH3YL1 (**Supplementary figure 4**). It has been shown that IL3RA (IL3 receptor alpha) is highly expressed on AML cells (Ehninger et al., 2014) and can be targeted using an anti CD123 monoclonal antibody in CML blast cells (Nievergall et al., 2014). Genes correlating negatively with CML progression included CSF2RA (GM-CSF apha receptor) which has been shown to be implicated in AML biology (Matsuura et al., 2012). ANPEP is the gene coding for CD13 antigen, which has been shown to be associated with monocytic / myeloid differentiation (Xu, 2006). It is of interest to note that we have identified in this screen, this gene to be negatively correlated with CML progression (**Supplementary figure 5**). The relevance of some of the genes identified in the progression of CML in our screen will require further study.

In summary, we report in this work the possibility of inducing a genomic instability at the level of a leukemic, patient-derived iPSC. The genomic alterations were detected by the use of CGH arrays and interestingly, upon hematopoietic differentiation, we were able to show cytological and several genomic features of blast crisis in this patient specific genomic background. It remains to be determined if the patients that we analyzed for this purpose was prone to the development of such a tool due to the initial resistance of the leukemic cells to Imatinib. It remains also to be determined if the stem cell model that we generated this way could lead to the generation of an in vivo model in immunodeficient mice. This first demonstration of the feasibility of using this approach to model blast crisis could open new perspectives for the future use of CML iPSC for disease modeling and eventually for identification of novel targets is they are validated in primary patient samples.

## FIGURE LEGENDS

**Supplementary figure 1:**
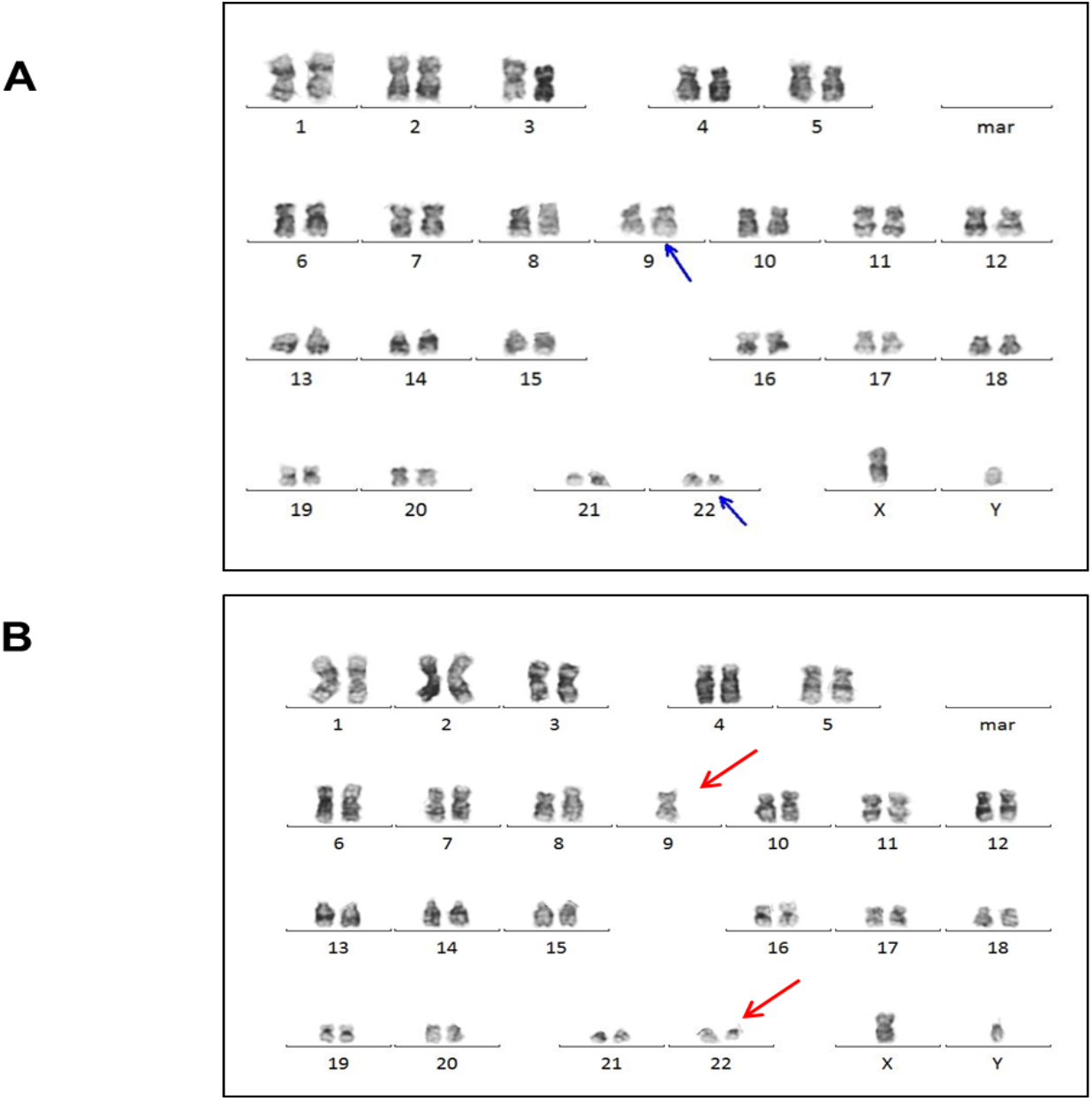
Evaluation of additional chromosome abnormalities in CML-iPSC after long term ENU culture. **(A)** Karyotype of CML IPSC with Ph1 chromosome and 9q+. **(B)** CML IPSC caryotype after 2 months of ENU exposure showing loss of Chromosome 9.

**Supplementary figure 2:**
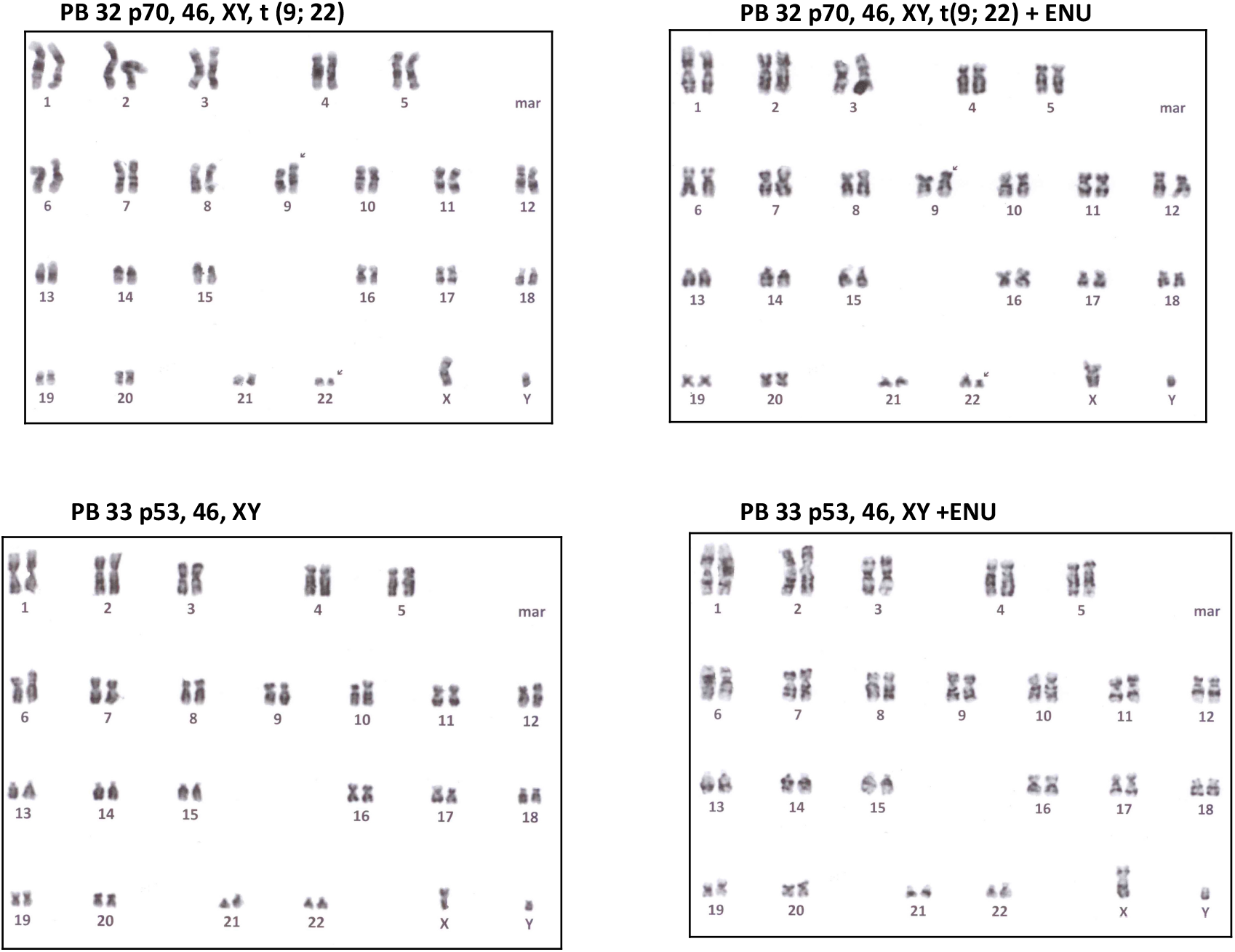
Representative cytogenetic analyses of PB32 (CML-IPC) and PB33 (control iPSC showing with and without long term ENU culture. **P53: passage 53, P46: passage 46.**

**Supplementary figure 3:**
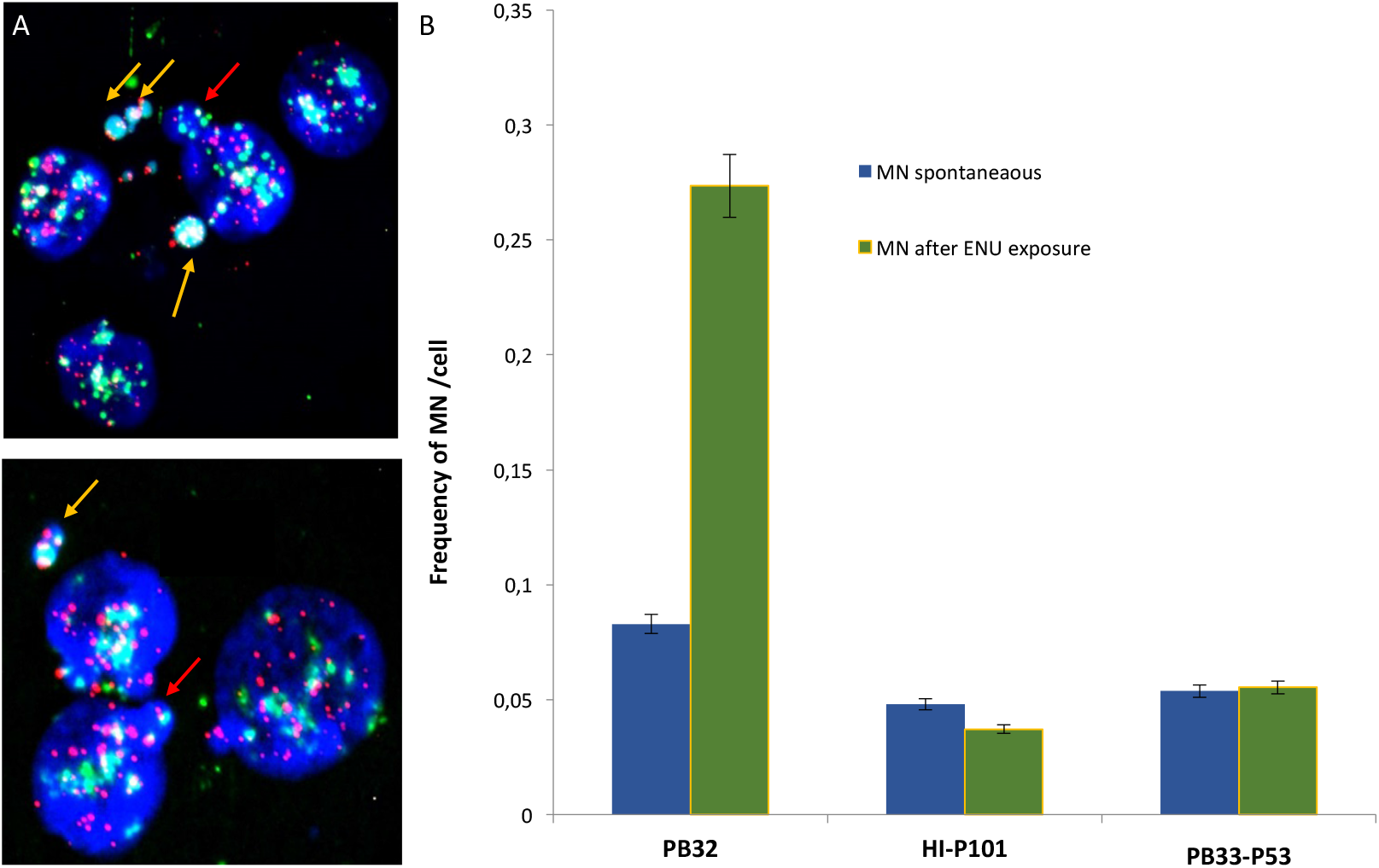
Evlauation of global **genomic instability by micronucleus assay in CML-IPSC, control iPSC and human embryonic stem cell line H1 cell line before and after ENU exposure.** **(A)** Photograph of CML-IPS colonies after ENU exposure. Yellow Arrow show micronucleus and white arrow shows nucleus bud (NBUD’s). **(B)** Micronucleus frequencies of CML-IPSC compared to control ESC and IPSC with and without long term ENU exposure.

**Supplementary figure 4:**
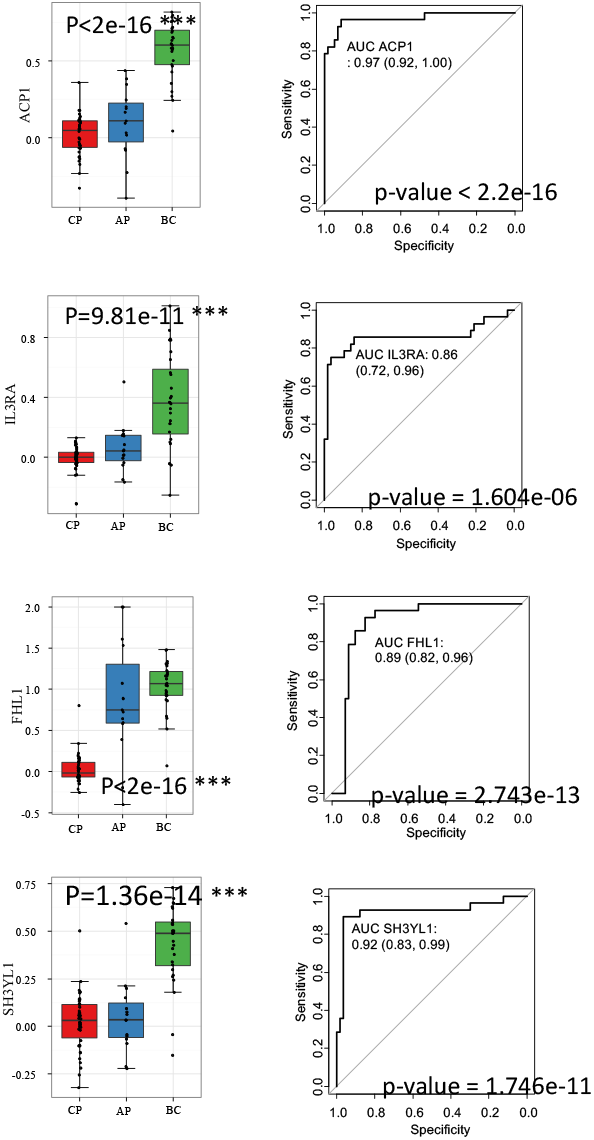
Genes with negative correlation to CML progression.

**Supplementary figure 5:**
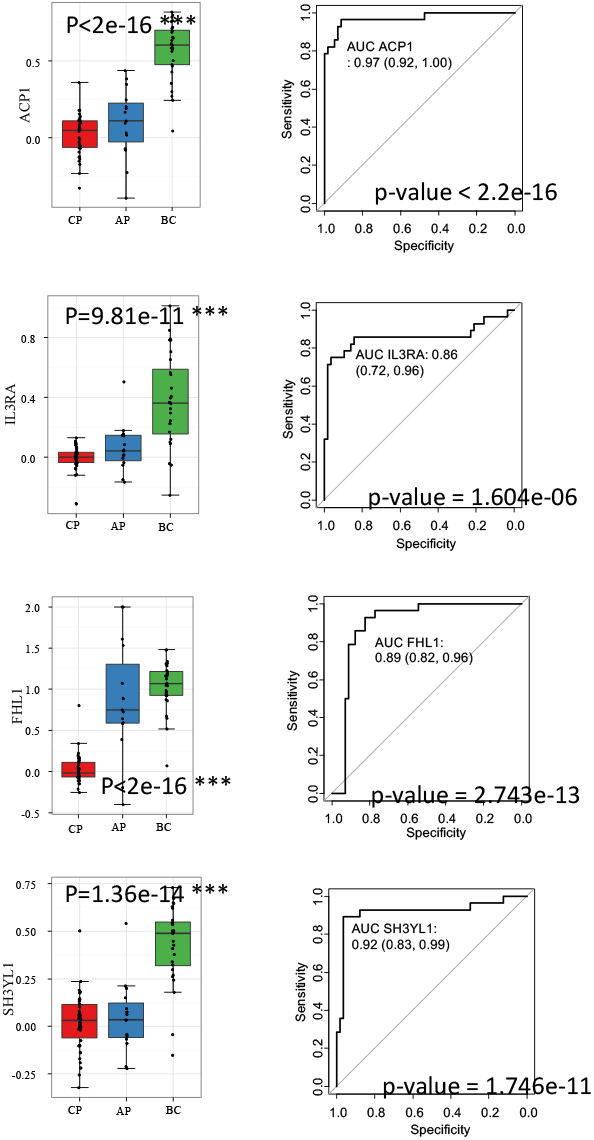
Genes with positive correlation to CML progression.

**Supplementary table 1**: **CNV without polymorphism from EB-derived CFCs.**

**Supplementary table 2**: **CNV without polymorphism from Bl-CFCs.**

**Supplementary table 3**: **Predictive genes.**

